# Survival cost sharing among altruistic full siblings in Mendelian population

**DOI:** 10.1101/2024.09.17.613452

**Authors:** József Garay, Inmaculada López, Zoltán Varga, Villő Csiszár, Tamás F. Móri

**Author notes:** Corresponding author: Inmaculada López. **Email:**.

## Abstract

**Background:** We focus on Haldane’s familial selection in monogamous families in a diploid population, where the survival probability of each sibling is determined by altruistic food sharing with its siblings during starvation. An autosomal recessive-dominant or intermediate allele pair uniquely determines the altruistic or selfish behavior, which are coded by homozygotes. We focus on the case when additive cost and benefit functions determine the survival probability of each full sibling.

**Results:** We provide conditions for the existence of the altruistic and selfish homozygote. We show that the condition of evolutionary stability of altruism depends on the genotype-phenotype mapping. Furthermore, if the offspring size increases then the condition of evolutionary stability of altruism becomes stricter. Contrary to that, for the evolutionary stability of selfish behavior it is enough if the classical Hamilton’s rule does not hold. Moreover, when the classical Hamilton’s rule holds and the condition of evolutionarily stability of altruism does not hold, then the selfish and altruistic phenotypes coexist.

**Conclusions:** In summary, the classical Hamilton’s rule is a sufficient condition for the existence of altruism, but it alone does not imply the evolutionary stability of the pure altruistic homozygote population when the altruistic siblings share the cost of altruism.

## Background

Haldane [1] was interested in “*familial selection*”, where the size of the family is strictly limited by the food resource, and more newborns are produced than can survive to enter the among the members of the same family (e.g. [2]). Contrary to Haldane, in the present paper we focus on the opposite situation, when within monogamous, and diploid families struggle for existence, by food sharing the siblings help each other to survive during a period when the food is limited ([3, 4]). Clearly, during starvation, food sharing is an altruistic behavior since it decreases the survival probability of the donor, while increases that of the recipient. Here we focus on the following case: Whenever a sib is starving, all altruistic sibs help the starving one to survive by donating the necessary food such that the total quantity of the donated food does not depend on the number of donors. Suppose the received food increases the survival of the starving sib by *b*. If there is only one altruistic sib, then it is the only to donate, and the donation reduces its own survival probability by *c*. If there are *n* altruistic sibs, then the survival probability of each will be reduced by *c*/*n*, since they share the costs. We emphasize that in each given food sharing event, we have an additive cost model ([4]). Here we will look for a condition of subsistence of sib altruism.

In Haldane’s familial selection model, full siblings (sharing both parents) interact, thus it clearly belongs to the aegis of kin selection, in the sense of Maynard Smith [5]. Kin selection theory focuses on the genetic relatedness between the recipient and the actor ([6, 1, 7]). The heuristic basic idea of the kin selection theory is the inclusive fitness effect: The altruistic gene can increase its evolutionary success when an individual having the altruistic gene promotes the survival of its sibling, who also carries the same altruistic gene, thus the altruistic individual indirectly increases the frequency of the altruistic gene. The well-known mathematical formulation of the idea of this indirect effect is the classical Hamilton’s rule ([7]): the altruism will spread if *rb* > *c*, where *b* is the fitness benefit to the recipient, *c* is the fitness cost to the actor and *r* is the coefficient of genetic relatedness between the interacting individuals. We also focus on altruistic helping with benefit *b* and cost *c*, but not in pairwise interactions. We emphasize that our game is not a standard evolutionary game, like e.g. the well-known Public Goods Game (e.g. [8]), in which the individuals’ benefit may be realized in fecundity, and all individuals have equal benefit from the altruistic act. Our game is a survival game (cf. [9]), where only one individual gets the benefit (sibling’s life is rescued by its altruistic siblings), and the costs of helping (decreasing altruistic siblings’ survival rate) are shared by the altruistic siblings (i.e. the altruistic siblings have no direct benefit). We concentrate on survival for two main reasons. Survival is a prerequisite for reproduction, thus helping in survival seems an important evolutionary strategy between siblings. Moreover, in our survival game, during an altruistic act there is no direct benefit for altruistic individuals (contrary to the public goods game); thus, only indirect fitness effects can promote altruism, i.e. rescue its parent’s altruistic allele. As a result, in the survival game considered here, the direct benefit of altruistic sibling cannot mask the indirect effect.

Over the last decade, the classical Hamilton’s rule has been a topic of controversy ([10-12]). Van Veelen made an effort to find the connection between kin and group selection theories, but he pointed out that these theories, in general, are not equivalent ([13-15], [10]). Furthermore, he was looking for conditions under which the classical Hamilton’s rule gives a right prediction. Based on replicator dynamics, he found that in a linear model (when the fitness effects are additive and homogenous in the number of altruistic siblings) the classical Hamilton’s rule is correct ([14, 16]). Although the replicator dynamics can handle asexual populations ([17]), contrary to van Veelen, we focus on diploid sexual populations.

The group selection method focuses on the advantage of group living ([18-20]]). Haldane’s familial selection is connected to the group selection too, since the interaction happens in family. There are ingredients of the group selection theory which are important in Haldane’s familial selection. The interaction is not well-mixed in the whole population and altruistic act can be a common action of full siblings ([4]). Usually, this interaction is given by a multi-player game ([21]) with the synergetic effect of the number of altruists on the individual fitness of the group (i.e. family) members. However, the other two ingredients of the group selection theory do not occur in Haldane’s familial selection. In group selection theory, there is a direct competition between different groups. Contrary to that, in Haldane’s familial selection there is no competition between different families. Moreover, in group selection theory, the groups are “quasi” stable and the phenotypic composition of the groups is determined by the formation process of the groups. Contrary to that, in Haldane’s familial selection, the survived newborns leave their family, and form new families according to the mating system. The phenotypic composition of the families is determined by the genetic system ([3, 4]) and the family’s phenotypic composition is fixed by the genotype of the parents. For the perspective of the present paper, the heuristic basic idea of the group selection theory is that the common action by group members (within groups and between groups) should be evolutionary advantageous for them. Here we only focus on Haldane’s familiar selection where there is common action within families, but no intra-familiar competitions.

We agree with Van Veelen and his coauthors, who pointed out these theories are not equivalent ([12], [22]). However, from the viewpoint of Haldane’s familial selection, the heuristic basic ideas of the above two theories are reasonable at the same time. On the one hand, the indirect fitness effect can work in diploid sexual families. On the other hand, the group effect (common action by altruistic siblings on the survival rate) can also work, because the selection takes place within a family. Since a direct combination of these theories is questionable ([12], [22]), in the framework of a population genetic model one can bring together the heuristic basic ideas of these theories ([3, 4, [23]]). Indeed, in monogamous diploid families, the genetic relatedness between full siblings is ½, thus altruistic help can increase the frequency of the altruistic gene indirectly. Furthermore, the survival probability of each sibling depends on its siblings’ common action (in other words, the group effect takes place). To make it clear what the role of the indirect fitness and group effects is, we must follow van Veelen and his coauthors ([24]), treating population genetic models rigorously.

The basis of our investigation is the orthodox Darwinian view, in the sense that we focus exclusively on the frequency change of genotypes in time. We think focusing on genotypes is the most direct and simplest way to study the evolution in the framework of Haldane’s familial selection. Why do we follow this orthodox Darwinian view? Below we mention some other possibly applicable methods and the reason why we think that they are not useful in Haldane’s familiar selection, but without a complete overview, of course.

The applicability of the Price equation was criticized by [12, 13, 25].

The allele frequency based model is depending on Haldane’s familial selection. The reason is the following. Under panmixia the embryos follow Hardy–Weinberg equilibrium, but during maturation time, the survival rate of each juvenile depends on the genotypes of their full siblings, which are determined by their parents’ genotypes. Consequently, the parental population is not in Hardy–Weinberg equilibrium. The main technical problem is that in Haldane’s familial selection model there is no bijection between genotype and phenotype distributions without genetic equilibrium [26]. In mathematical terms: the genotype distribution cannot be reconstructed unequivocally from a given allele distribution when the population is not in Hardy–Weinberg equilibrium. Thus, if the parental population is not in Hardy–Weinberg equilibrium, then an infinite number of genotype (phenotype) distributions correspond to the same allele distribution. The number of genotypes is always higher than the number of alleles. Consequently, according to the dominant-recessive inheritance of altruism, at the same parental allele distribution we have an infinite number of parental genotype distributions, and for different parental distributions the relative frequency of altruistic siblings is quite different in the whole population.^1^ The natural selection operates on the diploid level. Consequently, the natural selection takes place in a higher dimensional genotype space than the allele space. Thus, the consecutive populations are never in Hardy–Weinberg equilibrium; we can only follow the evolutionary change by following the frequency change of genotype distributions. We also mention that Allen and his co-workers [28-31] built up a general stochastic mathematical framework for natural selection. At each time step they calculated which alleles were replaced by copies of others as a result of interaction, reproduction, mating, and/or death. Contrary to that, in essence they also follow the genotype distribution change from generation to generation. Finally, in the population genetics models, when the juveniles’ survival rates depend on the genotypes of their parents, the state variable of the model must be the genotype distribution [32-36].

The applicability of individual-based fitness methods (e.g. [27]) is ambiguous in Haldane’s familial selection. In diploid sexual population, the parent population produces each juvenile genotype (see Table 1), since all genotypes produce other genotypes. For instance, in all families founded by two different homozygotes, all the offspring are heterozygotes. In other words, the principle of replicator dynamics, i.e. “*i*-type only from *i*-type”, does not hold when we focus on the frequency of genotypes. Moreover, an observer who can identify the different genotypes, can collect all information about natural selection. Thus in Haldane’s familial selection model the production of different genotypes is the key problem [23], and not the individual fitness (see genotype dynamics, Equation (4) later). We note, when strict reproduction occurs (i.e. *i*-type only produces *i*-type), the individual fitness is calculated as the quotient of the total production of the *i*-type and the total number of *i*-type individuals. Thus, practically, the production rate is a more basic notion than the average production rate (i.e. individual fitness).

**Table 1.** Genotype survival table based on the mating table, with phenotypes, and Mendelian probabilities *m*_*k*(*ij*)_.

| Genotypes<br>of parents | Average<br>number<br>of<br>couples | Average number of survived family members, for genotypes $G_1$, $G_2$, $G_3$ | | |
| --- | --- | --- | --- | --- |
| | | $G_1 \equiv ([a],[a])$<br>altruistic | $G_2 \equiv ([a],[A])$<br>mixed phenotype $p$ | $G_3 \equiv ([A],[A])$,<br>selfish |
| $G_1 \times G_1$ | $\frac{N}{2} x_1^2$ | $m_{1(11)} = 1$<br>$n_{1(11)} = n(a + b - c)$ | $m_{2(11)} = 0$<br>thus<br>$n_{2(11)} = 0$ | $m_{3(11)} = 0$<br>thus<br>$n_{3(11)} = 0$ |
| $G_1 \times G_2$ | $N x_1 x_2$ | $m_{1(12)} = \frac{1}{2}$<br>$n_{1(12)} =$<br>$\frac{n}{2}a + \frac{n}{2}\left(1 - \left(\frac{1-p}{2}\right)^{n-1}\right)b$<br>$-\frac{n}{1+p}\left(1 - \left(\frac{1-p}{2}\right)^{n-1}\right)c$ | $m_{2(12)} = \frac{1}{2}$<br>$n_{2(12)} =$<br>$\frac{n}{2}a + \frac{n}{2}\left(1 - \left(\frac{1-p}{2}\right)^{n-1}\right)b$<br>$-\frac{pn}{1+p}\left(1 - \left(\frac{1-p}{2}\right)^{n-1}\right)c$ | $m_{3(12)} = 0$<br>thus<br>$n_{3(12)} = 0$ |
| $G_1 \times G_3$ | $N x_1 x_3$ | $m_{1(13)} = 0$<br>thus<br>$n_{1(13)} = 0$ | $m_{2(13)} = 1$<br>$n_{2(13)} = na$<br>$+n(1 - (1-p)^{n-1})(b - c)$ | $m_{3(13)} = 0$<br>thus<br>$n_{3(13)} = 0$ |
| $G_2 \times G_2$ | $\frac{N}{2} x_2^2$ | $m_{1(22)} = \frac{1}{4}$<br>$n_{1(22)} =$<br>$\frac{n}{4}a + \frac{n}{4}\left(1 - \left(\frac{3-2p}{4}\right)^{n-1}\right)b$<br>$-\frac{n}{1+2p}\left(1 - \left(\frac{3-2p}{4}\right)^{n-1}\right)c$ | $m_{2(22)} = \frac{1}{2}$<br>$n_{2(22)} =$<br>$\frac{n}{2}a + \frac{n}{2}\left(1 - \left(\frac{3-2p}{4}\right)^{n-1}\right)b$<br>$-\frac{2pn}{1+2p}\left(1 - \left(\frac{3-2p}{4}\right)^{n-1}\right)c$ | $m_{3(22)} = \frac{1}{4}$<br>$n_{3(22)} =$<br>$\frac{n}{4}a + \frac{n}{4}\left(1 - \left(\frac{3-2p}{4}\right)^{n-1}\right)b$ |
| $G_2 \times G_3$ | $N x_2 x_3$ | $m_{1(23)} = 0$<br>thus<br>$n_{1(23)} = 0$ | $m_{2(23)} = \frac{1}{2}$<br>$n_{2(23)} =$<br>$\frac{n}{2}a + \frac{n}{2}\left(1 - \left(\frac{2-p}{2}\right)^{n-1}\right)b$<br>$-n\left(1 - \left(\frac{2-p}{2}\right)^{n-1}\right)c$ | $m_{3(23)} = \frac{1}{2}$<br>$n_{3(23)} =$<br>$\frac{n}{2}a + \frac{n}{2}\left(1 - \left(\frac{2-p}{2}\right)^{n-1}\right)b$ |
| $G_3 \times G_3$ | $\frac{N}{2} x_3^2$ | $m_{1(33)} = 0$<br>thus<br>$n_{1(33)} = 0$ | $m_{1(33)} = 0$<br>thus<br>$n_{2(33)} = 0$ | $m_{1(33)} = 1$<br>$n_{3(33)} = na$ |

The direct application of inclusive fitness is also depending on Haldane’s familial selection.

There are three main reasons for that.

1. In the framework of Haldane’s familial selection model, the interaction takes place in families, so the benefit and cost of siblings depend only on the genotypes of their parents. The phenotypic composition of each family does not depend on the frequencies of the different parent genotypes in the whole population. Thus, the sib’s benefit and the sib’s cost within families are independent of the genotype frequencies in the whole population, and hence they are “independent” of the allele frequencies in the whole population, too.
2. In the present-day theory of kin selection, in terms of general Hamilton’s rule, the parameters of benefit *b* and cost *c* are functions of the allele frequencies ([6]). In Haldane’s familial selection the average benefit/cost of the juvenile genotypes with respect to the whole population depends on the genotype distribution of the parental population.
3. Moreover, many parental genotype distributions have the same allele distribution. Thus, the allele distribution contains less information about the selection process than the genotype distribution.
4. The general definition of genetic relatedness says that it is the regression of the genotypes of social partners on the genotype of the focal individual ([37-39]). In the framework of Haldane’s familial selection under monogamy, the genetic relatedness is exactly ½ between interacting full siblings independently of the state of the whole population.
5. We are interested in the condition under which the rare mutant selfish allele cannot invade a resident altruistic homozygote population. When the mutant allele is rare enough, using the mean-field model, we are interested in what population-state-independent condition guarantees that the relative frequency of altruistic homozygotes always increases. In other words, we are looking for a condition that does not depend on the genotype (so on the allele) frequencies, like in the case of the classical Hamilton’s rule.

Of course, we do not claim that the above listed methods cannot give the same results as we get. We only mentioned some points why we follow the most direct, orthodox Darwinian method, i.e. we focus only on the change of genotype frequencies in the whole population.

Why do we focus on diploid populations? Firstly, since Haldane considered that. Furthermore, one may think the results in a haploid population are almost the same as the results in a diploid population. This is one of the possible reasons why authors mostly consider haploid models (see e.g. [6], [17]). There is an essential difference between the haploid and diploid model. Namely, each diploid individual has two alleles together determining the phenotype. In other words, in a haploid model there is no Mendelian inheritance, since a haploid individual has only one allele, thus there is no meaning of dominance between alleles. Thus if anyone is interested in the effect of Mendelian inheritance, she/he must consider diploid populations, as we have done.

In this paper, we focus on diploid, panmictic populations with Mendelian inheritance within monogamous families ([3, 4]). There are two main consequences of Mendelian inheritance in a diploid population: Firstly, the genotype determines the phenotype. Secondly, the phenotypic composition of each family is determined by the mating system, the genotypes of parents, and the genotype-phenotype mapping together. In general, the population genetics models predict that the fixation of the altruistic behavior depends on the genotype-phenotype mapping (e.g. [3, 40, 41, 42]). However, in pairwise interaction with additive cost and benefit functions, the population genetics models give the same results as Hamilton’s method ([3-4, 19]), but it does not in the non-additive situation ([43, 4, 14, 16]). We emphasize that these results are in harmony with the above recalled results by van Veelen and his coauthor, based on replicator dynamics ([14, 16, 17]).

In a family, not only pairwise interactions are possible. Since we can assume there are more than two siblings in families, we can consider non-pairwise interactions. Remember, in the standard synergetic group effect, the group members’ action is common (non-pairwise), which in a non-linear way determines the benefit of each member of the group ([4, 44]). In this paper, we also focus on a non-pairwise interaction, i.e. when full siblings (sharing both parents) share the cost of altruism. Moreover, an additive cost and benefit function determines the survival probability of each full sibling ([14]). Shortly, we consider additive and non-pairwise interactions. We emphasize that our model is additive in the sense that both the costs and benefits sum up, but by cost sharing the homogeneity is violated. Therefore, our model is not linear.

Now the main question of this paper arises: What is the role of the classical Hamilton’s rule in diploid Mendelian population with cost sharing among altruistic siblings?

## Methods: Model for cost sharing

We will use the already introduced mating table based population genetic model ([3]) for a diploid species with internal fertilization. For the reader’s convenience, we present an overview of this model setup.

For the sake of simplicity, we only consider one autosomal locus with two alleles *a* and *A*. Since here the survival rate of each juvenile depends on its parents’ genotypes, the diploid parental population is not in Hardy–Weinberg equilibrium. Consequently ([45]), the state variables of our model must be the frequencies of genotypes. Let *x* = (*x*_1_, *x*_2_, *x*_3_) be the frequency vector of genotypes *G*_1_ = ([*a*],[*a*]), *G*_2_ = ([*a*],[*A*]) and *G*_3_ = ([*A*],[*A*]) in the present generation. Then

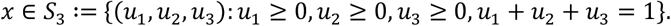

### Genotype-phenotype mapping

We assume that the genotypes uniquely determine the phenotypes. The homozygotes *G*_1_ = ([*a*], [*a*]) and *G*_3_ = ([*A*], [*A*]) are altruistic and selfish, resp., and heterozygote *G*_2_ = ([*a*], [*A*]) is either altruistic or selfish. We will analyze the following general setup. The phenotype of a heterozygote individual is altruistic with probability *p* ∈ [0, 1], and selfish with the complementary probability 1 − *p*. Particularly, altruism is recessive, dominant, or intermediate (additive), according to *p* = 0, 1, or 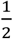. We note that *p* gives the *degree of dominance* of the altruistic allele.

### Structure of the population

The species is monogamous, thus relatedness between full siblings is ½. The population size *N* is very large, and we consider *panmixia*, so *N*/2 couples are formed at random. We note that the random mating excludes inbreeding, since the population is large enough. Formally, at the population state *x* ∈ *S*_3_, the relative frequency of the mating pair with genotypes *G*_i_ and *G*_*j*_ is equal to *x*_*i*_*x*_*j*_. Moreover, the parents’ genotypes, according to the Mendelian inheritance, determine the genotype of their offspring. Denote by *m*_*k*(*ij*)_ the probability that a pair with *G*_*i*_ and *G*_*j*_ genotypes will have an offspring with genotype *G*_*k*_ (see Table 1). Since the genotype determines the phenotype, the phenotypic composition of the family of parents with *G*_i_ and *G*_j_ genotypes follows either binomial or multinomial distribution. Moreover, the relative frequencies of different families depend on the relative frequencies of the parents. In summary, the population structure depends on the mating system, the parents’ genotype distribution *x*, and the genotype-phenotype mapping at the same time. In our *n*-person *altruistic interaction* there are exactly *n* siblings in each family.

### Altruistic interaction with cost sharing

By food sharing siblings can help each other to survive during an extreme food-poor period. For surviving a period of starvation, only a given quantity of food is needed. The main point here is that, under a series of predator attacks each individual must survive all attacks (so the product of survival probabilities is the final survival rate) [9]. Contrary to that, in starvation each individual must survive a period, therefore here the additive model seems more reasonable than the multiplicative one. Thus, we can consider an additive model for survival. Each sibling having at least one altruistic sib receives benefit *b* altogether, which is collected from its altruistic sibs equally. Thus, in a family with *k* altruistic and *n* − *k* selfish siblings, the payoff of each altruist is equal to *a* − (*n* − 1)*c*, if *k* = 1 (where *a* is the baseline survival rate without any extra food from siblings) and to 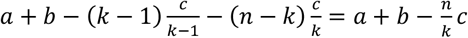, if *k* ≥ 2. At the same time, each selfish individual’s payoff is *a* + *b*, if *k* ≥ 1, and *a*, if *k* = 0. More precisely, the survival probability function has to be truncated at 0 and 1, that is, if our formula gives the value *p*, the survival probability is max{min{1, *p*}, 0}. However, if a single altruistic act has only a relatively small effect on the survival probability of the helper and the beneficiary, then truncation is needed with but a practically negligible probability. For the sake of computational simplicity, we will omit truncation, that is, *c* ≤ *a*/(*n* − 1) and *b* ≤ 1 − *a* is assumed. This assumption resembles the usual weak selection, where all individuals perform approximately equal fitness [46, 47]. However, in our model we focus on the survival probability of different types, and we do not need that those are approximately equal. For instance, in our model the different genotypes may have survival rates 3/4, 1/2 and 1/4. Also observe that each offspring’s survival probability only depends on the phenotype of its siblings. In other words, the interaction is well-mixed in the families, but it is not well-mixed with respect to the whole population. We denote by *n*_k(ij)_ the average number of surviving sibs with genotype *G*_k_ in a family founded by parents with genotypes *G*_*i*_ and *G*_*j*_, and we compute all numbers *n*_*k(ij)*_ in SI-A.

In Table 1, we present the average survival rate of each genotype in different families. If there are altruistic ones among the siblings, then each sibling (wether altruistic or selfish) receives an additional *b* to its survival probability. The cost of this increase is equal to *c*, which is shared among the altruistic siblings equally. So each beneficiary contributes *b* − *c* to the cumulative payoff. If *k* − 1, there are only *n* − 1 beneficiaries, while for *k* > 1 all *n* siblings are beneficiaries.

Based on Table 1, the total number of individuals of genotypes *G*_1_, *G*_2_ and *G*_3_ in the next generation can be given as follows:

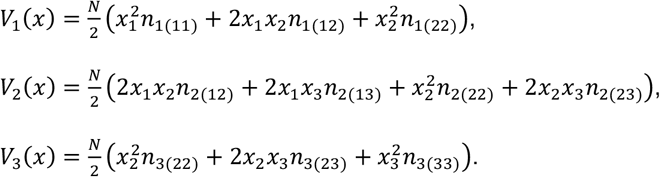

Observe that now the basic assumption of replicator dynamics is not valid, since here the *i*-th genotype is not necessarily born from *i*-th genotype. To emphasize this difference, we will call *V*_*i*_(*x*) *the production of the i-th genotype*. We will also need the notion of *total production* of the whole population, namely, 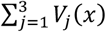.

## Results

### Static characterization of evolutionary stability

In this section, we are interested in the conditions under which the altruistic homozygote will make an *evolutionarily stable genotype distribution* (ESGD). We assume that the mutation is rare enough, as in the standard assumption of evolutionary game theory ([48]). Following [49], we will say that a homozygote is an ESGD if a rare mutant genotype cannot invade the resident homozygote population ([3]). Mathematically, this means that the relative frequency of the resident homozygote increases from generation to generation, provided mutation is sufficiently rare. Now, we recall the formal definition ([3]): The altruistic homozygote *G*_1_ is an from a neighborhood of *x*^*^ = (1,0,0) we have

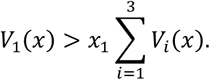

Firstly, we calculate the condition for the evolutionary stability of altruism. Applying a general result (see Theorem 1, [3]), we find the following (for mathematical details see Appendix A). If *p* < 1, then the pure homozygote altruistic population is an ESGD provided *n*_1(11)_ > 2*n*_2(12)_, which reads as

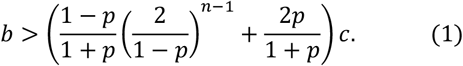

Particularly, if *p* = 0, the sufficient condition for the recessive altruism to be evolutionarily stable is

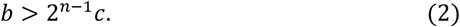

If *p* = 1 (altruism is dominant), then *n*_1(11)_ = 2*n*_2(12)_, therefore we need to check the second order conditions of the ESGD (Theorem 1, [3]). In our particular case, the general condition takes the following simpler form: genotype *G*_1_ is evolutionarily stable if

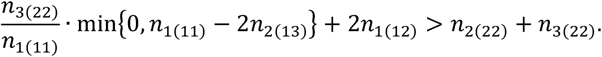

After some calculus we arrive at the simple condition

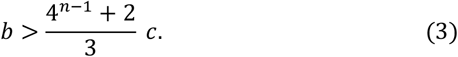

It is interesting that this condition coincides with that of the additive case, i.e., substituting *p* = 1/2 in Equation (1).

Now we are in the position to see the effect of genotype-phenotype mapping and that of cost sharing on the subsistence of altruism in Mendelian, diploid populations (see Table 2).

**Table 2.**
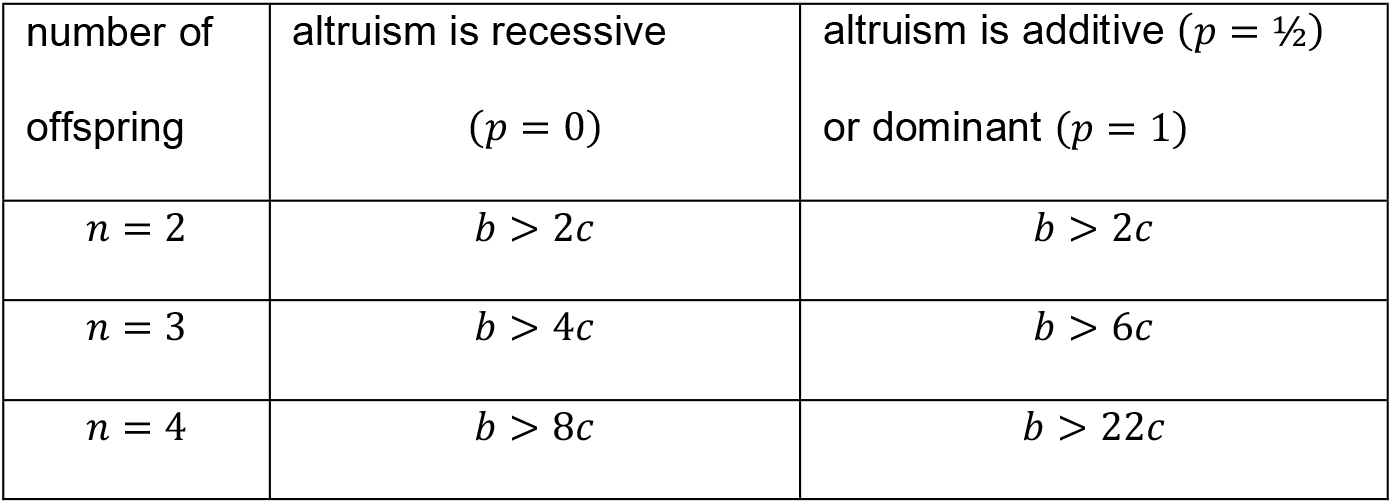
Sufficient conditions for altruism to be evolutionarily stable.

Now the survival probability is an additive function, and the relatedness between full siblings is ½ in each family. Thus, according to the state of art of the earlier results on altruism based on replicator dynamics (see references in the Introduction), one may expect that the classical Hamilton’s rule holds here. On the contrary, in the population genetics model for diploid species, we found this expectation true only when the offspring size was exactly two (see Table 2, *n* = 2), where cost sharing was not possible. We note that when the interaction is pairwise, the population genetics model pointed out that the classical Hamilton’s rule implied the evolutionarily stability of altruism ([3]).

We found that the genotype-phenotype mapping influences the condition of evolutionary stability of altruism (see Table 2). We think that the reason for this comes from the heuristic basic idea of group selection theory. The genotype-phenotype mapping determines the phenotypic composition of the families and the phenotypic distribution determines the number *s* of altruistic siblings, which determines the survival probability of each family member. Furthermore, we particularly found that when the altruistic allele is recessive, the conditions are less strict for arbitrary offspring size (see Table 2). The intuitive reason for this is that under recessive inheritance, at every mixed population state (*x* ≠ 1) there are less families having altruistic sibling, thus the selfish phenotype has less possibility for making use of altruistic siblings’ service.

Fixing the benefit *b*, we find that for the evolutionary stability of altruism, the cost must decrease as the family size *n* increases, assuming cost sharing among altruistic siblings (see Table 2). In other words, cost sharing implies stricter conditions for subsistence of altruism than the classical Hamilton’s rule. Thus, the naïve hypotheses that cost sharing could help in the fixation of altruism is wrong. This observation is rooted in the following two intuitive reasons: On the one hand, in the stochastic feature of Mendelian inheritance: when there is only one altruistic sib in a family, then it helps *n* − 1 selfish siblings, thus its cost is high. On the other hand, there are less altruistic events under cost sharing than in pairwise interactions: e.g., when there are three altruistic siblings, then each receives *b* by cost sharing, but 2*b* in pairwise interactions.

Secondly, we are also interested what conditions make the indirect net effect of the altruistic interaction remunerative for the altruistic gene [a] (see SI-B). We found that the classical Hamilton’s rule makes the altruistic interaction remunerative for the altruistic gene [a], under all considered genotype-phenotype mappings (i.e., for all *p* ∈ [0,1]), and at all population states (i.e. *x* ∈ *S*_3_). Observe that when the altruistic siblings can share the cost, the fact that the interaction is remunerative for the altruistic allele is not sufficient for the altruism to be evolutionarily stable (see Table 2). The reason for this is as follows. If we only concentrate on the fact that, on average, the altruistic interaction can increase the number of altruistic alleles (cf. the classical Hamilton’s rule), then we do not take account of the effect the altruistic interaction has on the number of selfish alleles. It can happen that the altruistic interaction increases the number of selfish alleles much more than that of the altruistic ones.

Thirdly, we found that if *b* < 2*c*, then in all here considered genotype-phenotype mappings, the selfish homozygote is an ESGD (see SI-A). This implies that, if the classical Hamilton’s rule holds, the altruistic gene spreads in the monomorphic selfish population. We emphasize that the stability condition of the selfish homozygote depends neither on the size of the family nor on the genotype-phenotype mapping.

Finally, since the mathematical conditions of ESGD of altruism and selfishness contradict each other, there is no bi stability here.

In summary, when there are more than two siblings sharing the cost of altruism (i.e., *n* ≥ 3), the classical Hamilton’s rule is a sufficient condition for the existence of altruism, but it does not imply the evolutionarily stability of the altruistic homozygote monomorphic population.

### Dynamical characterization of ESGD-s

The condition of classical evolutionary stability is a *local* one, since it rules out that any possible but rare mutant can invade the monomorphic homozygote population. So the following natural question arises: In the case of Haldane’s familial selection, what dynamics describes the natural selection *globally*? In [3] we introduced the *genotype dynamics*

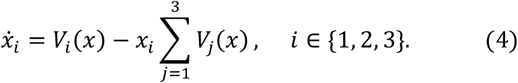

Observe that the genotype dynamics is based on the Darwinian tenet: The frequency of the *i*-th genotype increases in the next generation, if its production (i.e. *V*_*i*_(*x*)) is greater than its share would be in the total production of the whole population, corresponding to its proportion (i.e. 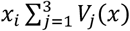). The following statement was proved (Theorem 2 in [3], SI C): If the altruistic homozygote *G*_1_ is an ESGD, then *x*^*^ = (1,0,0) is a locally asymptotically stable fixed point of the genotype dynamics, Equation (4). Clearly, the simplex of the genotype distributions is positively invariant with respect to the genotype dynamics, and the homozygote states are the only possible fixed points of the genotype dynamics on the border of the simplex of genotype distributions.

Here, we will use genotype dynamics, Equation (4) to gain some insight into the global behavior of our model. We are interested in what happens if the classical Hamilton’s rule is satisfied, and what happens if it is not. For this purpose, we consider three different genotype-phenotype mappings (altruism is either recessive, dominant, or intermediate). Concerning each genotype-phenotype mapping considered here, we focus on the following three types of numerical examples: 1. The altruistic homozygote is an ESGD. 2. Neither homozygotes are ESGD. 3. The selfish homozygote is an ESGD. We are interested in the cost sharing among siblings, therefore we consider *n* = 4, since cost sharing can only happen when the offspring size is greater than 2. For a unified representation, in all Figures 1, 2, 3 of the following illustrations, the same points of the simplex serve as the initial states of the trajectories: (0.86, 0.07, 0.07), (0.07, 0.86, 0.07), (0.2, 0.3, 0.5), (0.07, 0.07, 0.86), (0.48, 0.04, 0.48), (0.48, 0.48, 0.04) and (0.04, 0.48, 0.48):

**Figure 1.**
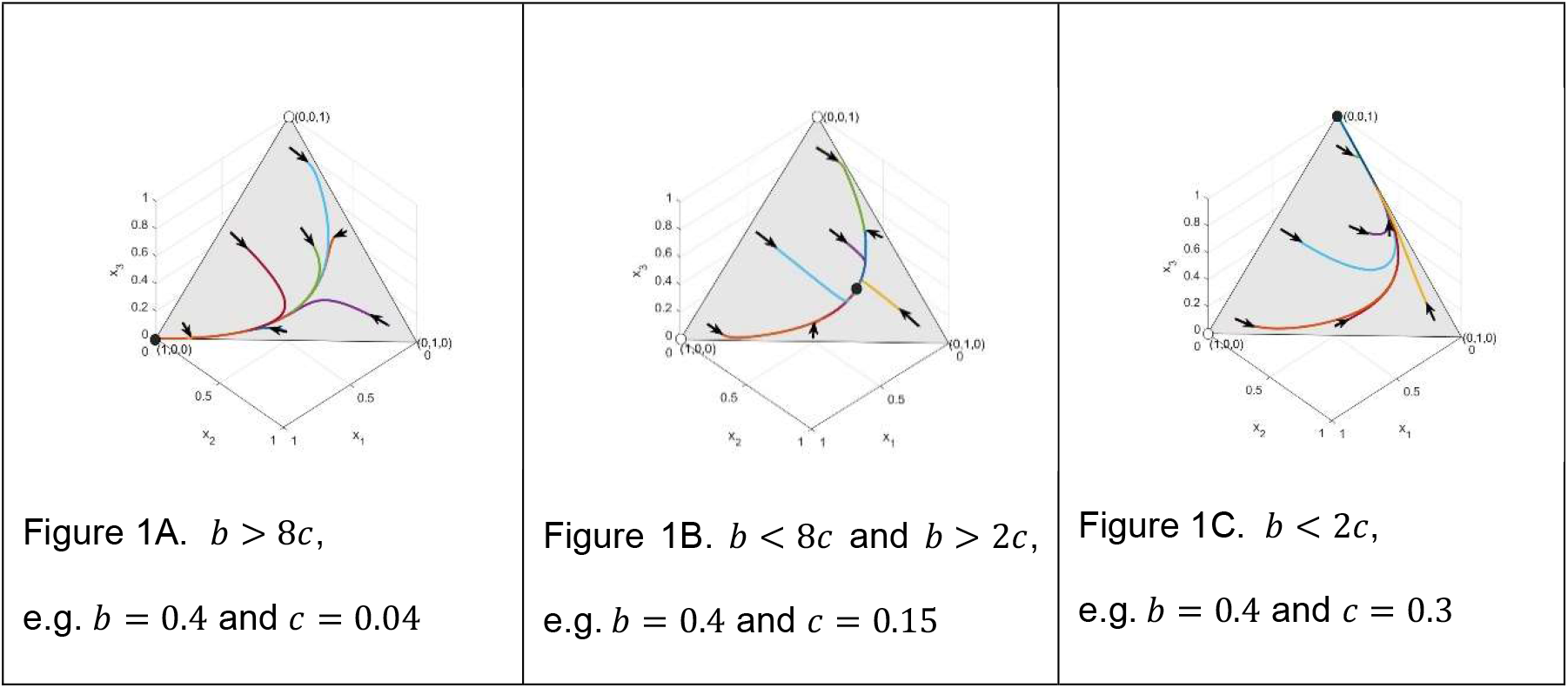
Firstly, we focus on the case when the altruistic behavior is recessive, i.e. *p* = 0. Figure 1A shows that, being pure ESGD, the recessive altruistic homozygote population is globally asymptotically stable. Figure 1B shows the case where neither of the homozygote populations are ESGD. Then there is a unique globally asymptotically stable equilibrium of the genotype dynamics, (0.2208, 0.5360, 0.2432). Figure 1C shows a case where Hamilton’s rule does not hold, and the pure selfish homozygote population is globally asymptotically stable.

**Figure 2.**
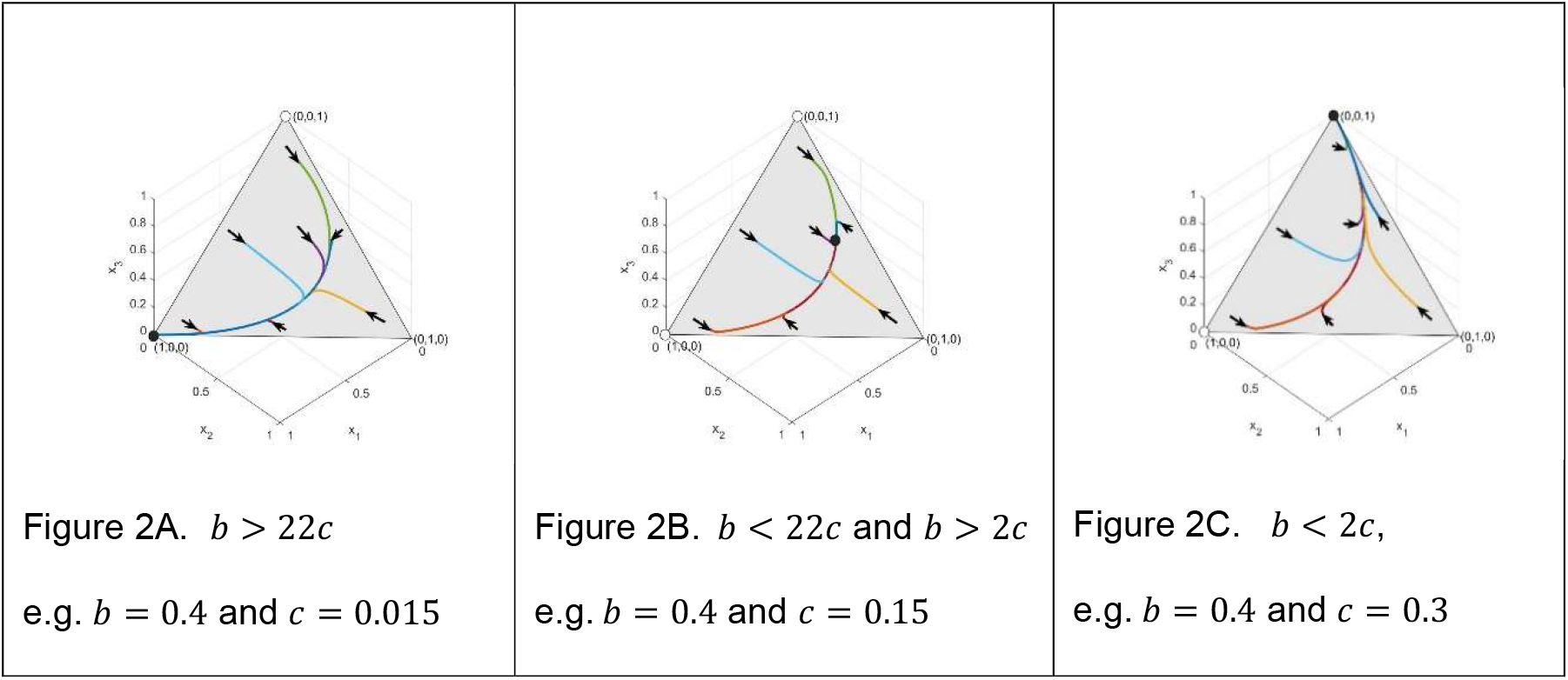
Secondly, we focus on the case where the altruistic behavior is dominant, i.e. *p* = 1, and *n* = 4. Figure 2A shows that, being pure ESGD, the dominant altruistic homozygote population is globally asymptotically stable. Figure 2B shows that when neither of the homozygote populations are ESGD, then there is a unique globally stable equilibrium, (0.1280, 0.4407, 0.4313) of the genotype dynamics. Figure 2C shows that when Hamilton’s rule does not hold, then the pure selfish homozygote population is globally asymptotically stable.

**Figure 3.**
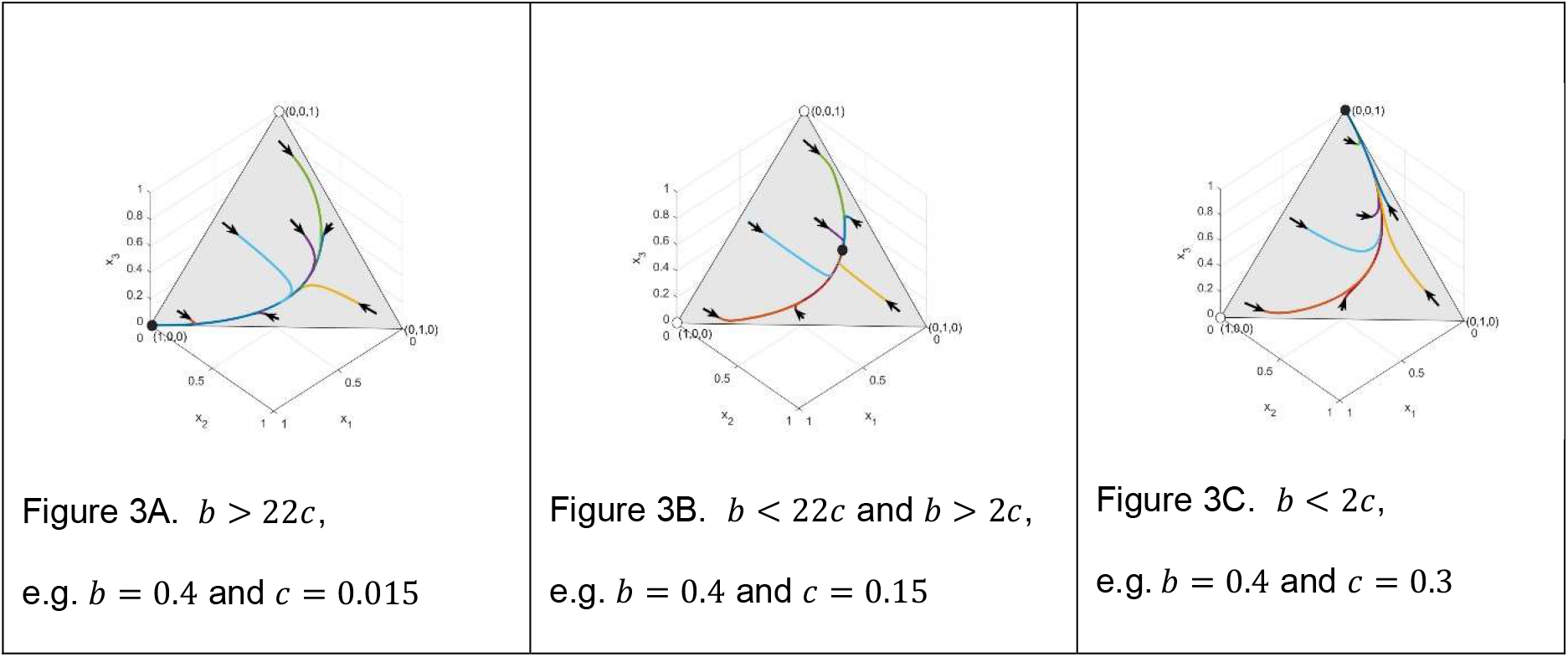
Thirdly, we focus on the case when the altruistic behavior is intermediate, i.e., 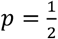, and *n* = 4. Figure 3A shows that the intermediate altruistic homozygote population, being a pure ESGD, is globally asymptotically stable. Figure 3B shows that when neither of the homozygote populations are ESGD, then there is a unique globally stable equilibrium of the genotype dynamics, (0.1603, 0.4828, 0.3569). Figure 3C shows that when Hamilton’s rule does not hold, then the pure selfish homozygote population is globally asymptotically stable.

## Discussion

Now we are in the position to summarize our results. In Results section, we already gave a condition for the altruistic homozygote to be an ESGD, which implies the local asymptotic stability of the pure altruistic homozygote population. Using genotype dynamics, Equation (4), we illustrated that for all here considered genotype-phenotype mappings, the pure altruistic homozygote population is globally asymptotically stable if the altruistic homozygote is an ESGD (see Figures 1A, 2A and 3A).

By using genotype dynamics, Equation (4), we illustrated that if the altruistic and selfish homozygotes are not ESGDs, then in all here considered genotype-phenotype mappings, the altruistic and selfish phenotypes coexist and there is a unique globally asymptotically stable polymorphic state (see Figures 1B, 2B and 3B). However, these fixed points are different according to the genotype-phenotype mapping.

Furthermore, we illustrated that if the classical Hamilton’s rule does not hold, then the pure selfish homozygote population is globally asymptotically stable (see Figures 1C, 2C and 3C).

Based on our dynamical studies, in all considered selection situations, it seems that the local stability properties of the homozygotes have an effect on the global behavior of the genotype dynamics.

## Conclusions

Haldane’s familial selection model includes heuristic ideas of kin and group selections each at a time.

Concerning the indirect fitness effect of altruism, we found the following:

a. The naive hypothesis, that cost sharing could help the fixation of altruism, is based on the fact that the survival probabilities of altruistic siblings decrease less. Contrary to that, we have found that if the offspring size increases then the condition of evolutionarily stability of altruism becomes stricter (see Table 2).
b. The genotype-phenotype mapping has an effect on the condition of the evolutionarily stability of altruism. Namely, if altruism is recessive, then it will fix under weaker conditions. The intuitive reason for this is that, under recessive inheritance, the average number of altruistic siblings in the whole population is less, thus the selfish phenotype has less possibility for making use of altruistic siblings’ service.
c. Contrary to point b), we found a universal condition for the evolutionary stability of selfish behavior (for all here considered genotype-phenotype mappings and arbitrary size of offspring). If the classical Hamilton’s rule does not hold, then the selfish behavior is an ESGD. The intuitive reason for this is that the classical Hamilton’s rule guarantees that the net indirect effect of the altruistic interaction is remunerative for the altruist gene. Observe that this intuitive reasoning comes from the kin selection theory.
d. In all here considered genotype-phenotype mappings, the classical Hamilton’s rule implies the stability of altruism if and only if there are only two siblings in each family, so when the interaction is pairwise, as cost sharing is not possible.
e. Moreover, when the classical Hamilton’s rule holds and the condition of evolutionarily stability of altruism does not, then the selfish and altruistic phenotypes coexist.

In summary, when the altruistic siblings share the cost, the classical Hamilton’s rule is a sufficient condition for the existence of altruism, but it does not imply the evolutionary stability of the altruistic homozygote.

Group living opens up the opportunity for common action of more than two individuals. Here we focused on the cost sharing among altruistic siblings. We have found that the group living advantage takes place in Haldane’s familial selection model, since we have shown the following.

1. The conditions for the evolutionarily stability of altruism are sensitive to the offspring size (see inequalities (1–3)).
2. Our observation that the genotype-phenotype mapping modifies the condition of evolutionary stability of altruism, came true due to the heuristic basic idea of group selection, since the phenotypic composition of each family is determined by this mapping.

In summary, group living (by common action) has an effect on the evolutionary stability of altruism under cost sharing among siblings, and modifies the classical Hamilton’s rule.

The inclusive fitness effect and the group effect together modify the conditions of existence of altruism in diploid populations. Thus, it is safe to say, that the population genetics model can at the same time handle the heuristic basic ideas of kin theory (i.e. the indirect fitness effect) and group theory (the common action of the group members takes an effect on the survival probabilities), but it does not combine the methods and/or models of these theories.

Finally, we point out one of our main observations. Based on the standard replicator dynamics ([50]) in asexual populations, Van Veelen ([14, 16]) found that when the individuals’ fitness depends linearly on the costs and benefits, then Hamilton’s rule implies the evolutionarily stability of altruism. The linear specification needs additivity and homogeneity in the number of altruistic siblings. In the case of cost sharing the homogeneity condition is not satisfied. This makes it likely that the homogeneity condition might be an important condition for Hamilton’s rule in other cases, too.

## Declarations

### Ethics approval and consent to participate

Not applicable.

### Consent for publication

Not applicable.

### Availability of data and materials

Not applicable.

### Author Contributions

Conceptualization: József Garay; Methodology: József Garay, Tamás F. Móri; Formal analysis and Investigation: Villő Csiszár, Tamás F. Móri; Visualization and numerical investigation: Inmaculada López; Writing - original draft preparation: József Garay, Inmaculada López, Zoltán Varga, Villő Csiszár, Tamás F. Móri; Writing - review and editing: József Garay, Inmaculada López, Zoltán Varga, Villő Csiszár, Tamás F. Móri; Supervision: József Garay.

## Acknowledgements

We thank Ádám Kun, Andy Gardner and Tibor Standovár for their valuable comments on the earlier version of this paper.

## Supporting Information

Since this is a purely theoretical work, we do not deal with data. As for the method, our study is built upon the concept of evolutionary stability, adapted to a population model based on the general mating table. Our results were obtained using mathematical analysis and differential equations. The numerical simulations were carried out in MATLAB R2023a environment. The work can be repeated following the calculations of the SI.

### SI-A: Static condition for evolutionarily stability

#### Phenotypic group determines payoff

Each sibling may have one of two phenotypes, altruistic and selfish, and this phenotype is genetically fixed. Homozygotes *G*_1_ = ([*a*], [*a*])and *G*_3_ = ([*A*], [*A*]) are altruist and selfish, resp., and heterozygote *G*_2_ = ([*a*], [*A*]) is either altruistic or selfish. We will analyze the following general setup. The phenotype of a heterozygote individual is altruistic with probability *p* ∈ [0, 1], and selfish with the complementary probability 1 − *p*. Particularly, altruist is recessive, dominant, or additive, according that *p* = 0, 1, or 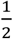. In our *n*-person donation game there are exactly *n* siblings in each family.

#### Cost sharing

Each sibling having at least one altruistic sib receives benefit *b* altogether, which is collected from its altruistic sibs equally. Thus, in a family with *k* altruistic and *n* − *k* selfish siblings the payoff of each altruist is equal to *a* − (*n* − 1)*c*, if *k* = 1, and to 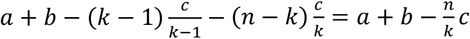, if *k* ≥ 2. At the same time, each selfish individual’s payoff is *a* + *b*, if *k* ≥ 1, and *a*, if *k* = 0. For the sake of simplicity, *c* ≤ *a*/(*n* − 1) and *b* ≤ 1 − *a* is assumed.

Let *n*_*i*(*jk*)_ denote the expected number of (geno)type *i* offspring in a family of type *G*_*j*_ × *G*_*k*_, *i, j, k* ∈ {1,2,3}. In what follows we compute these quantities. Throughout, 𝕀(.) will denote the indicator of the random event in brackets.

Let *X*_*cc*_, *X*_*hc*_, *X*_*hd*_, *X*_*dd*_ denote the (random) number of homozygote altruistic, heterozygote altruistic, heterozygote selfish, and homozygote selfish offspring, resp., in the family under consideration. Their joint distribution is multinomial (permitting degenerate ones with one or more parameters being zero).

For each family type,

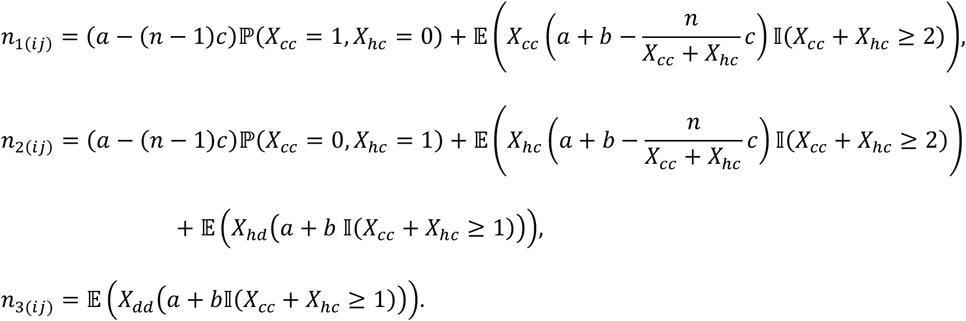

Suppose the joint distribution of (*X*_cc_, *X*_*hc*_, *X*_*hd*_, *X*_*dd*_) is general multinomial *M*(*n, r*_1_, *r*_2_, *r*_3_, *r*_4_). Then

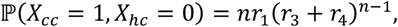

and the conditional distribution of *X*_*cc*_ given *X*_*cc*_ + *X*_*hc*_ = *k* is binomial 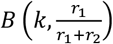, hence

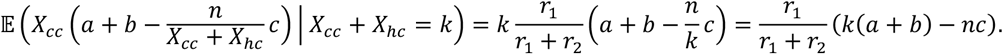

The distribution of *X*_*cc*_ + *X*_*hc*_ is binomial *B*(*n, r*_1_ + *r*_2_), thus

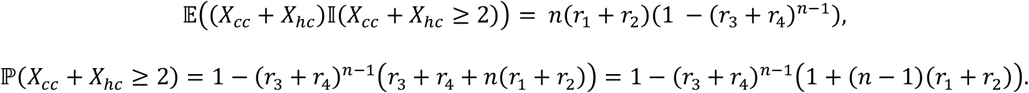

From all these it follows that

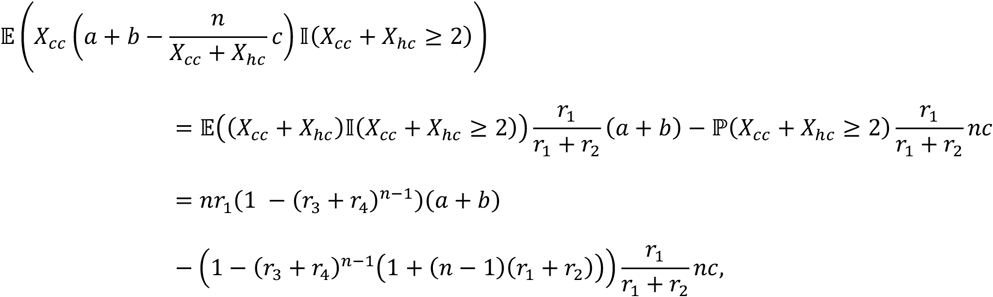

and consequently,

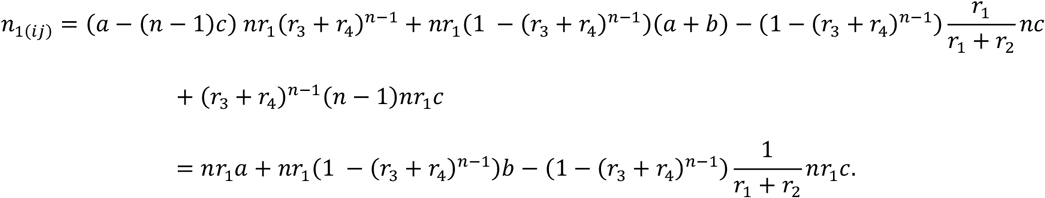

Let us turn to *n*_2(*ij*)_. As we have seen above,

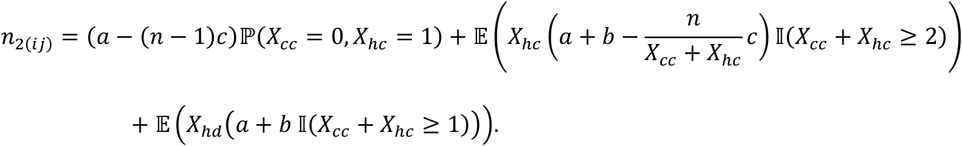

By analogy, the sum of the first two terms in the right-hand side is equal to

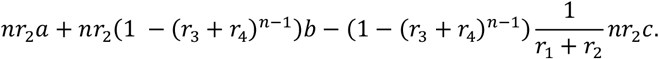

The last term can be treated as follows.

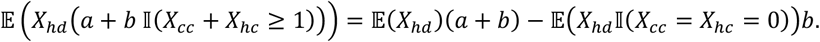

Here *X*_*hd*_ is binomial *B*(*n, r*_3_), hence 𝔼(*X*_*hd*_) = *nr*_3_.Moreover,

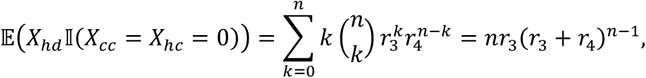

Thus

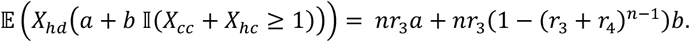

Consequently,

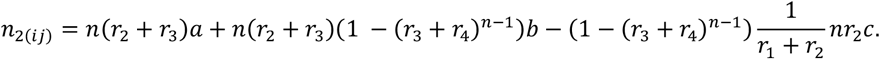

Finally, *n*_3(*ij*)_ can be obtained from the third term of *n*_2(*ij*)_ by interchanging *r*_3_ and *r*_4_, thus

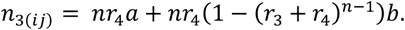

Let us specialize our results for each possible family types. In each cases we only have to find the probability distribution (*r*_1_, *r*_2_, *r*_3_, *r*_4_), and plug it into the formulas above.

**Family type** *G*_1_ × *G*_1_. Then *r*_1_ = 1, *r*_2_ = *r*_3_ = *r*_4_ = 0, thus

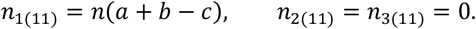

**Family type** *G* × *G*. Then 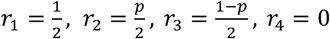. Consequently,

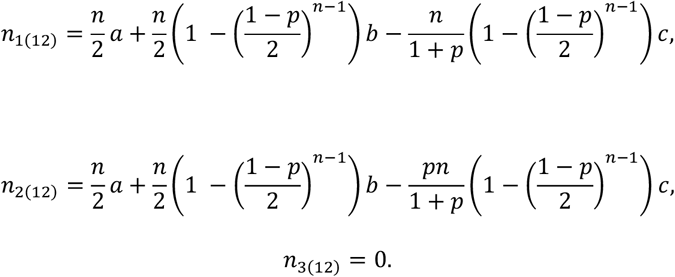

**Family type** *G*_1_ × *G*_3_. Then *r*_1_ = 0, *r*_2_ = *p, r*_3_ = 1 − *p, r*_4_ = 0. Hence

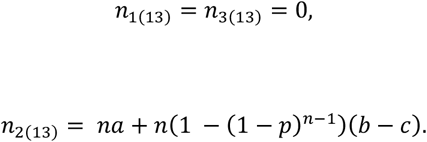

**Family type** *G* × *G*. Then 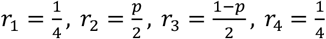, therefore

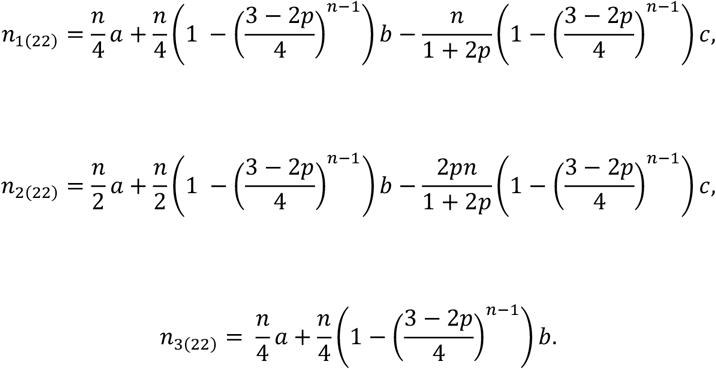

**Family type** *G* × *G*. Then 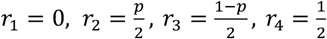. Thus, *n*_1(23)_=0

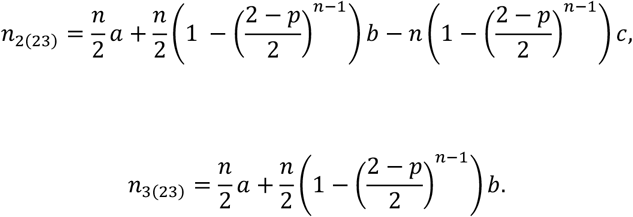

**Family type** *G*_3_ × *G*_3_. Then *r*_1_ = *r*_2_ = *r*_3_ = 0, *r*_4_ = 1, consequently

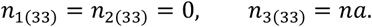

#### When is genotype *G*_1_ evolutionarily stable?

Let the pure altruistic genotype *G*_1_ be resident. According to [1], it is evolutionarily stable, if *n*_1(11)_ > 2*n*_2(12)_, that is,

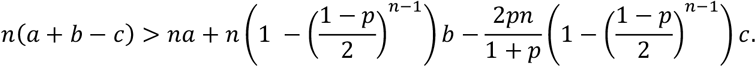

If *p* < 1, this means that

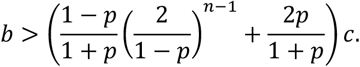

If *p* = 1 (altruism is dominant), then *n*_1(11)_ = 2*n*_2(12)_, therefore we need to check the second order conditions of the ESGD (Theorem 1, Garay et al., 2019). In our particular case, where there is only one primary and one secondary mutant phenotype (*G*_2_ and *G*_3_, resp.), the general condition puts on the following simpler form: genotype *G*_1_ is evolutionarily stable, if

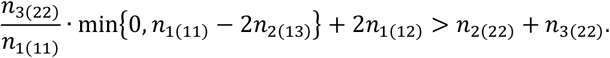

After some calculus we arrive at the condition

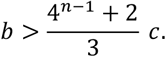

It is interesting that this condition coincides with that of the additive (*p* = 1/2) case. In the recessive case the sufficient condition for altruism to be evolutionarily stable is

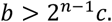

Note that for *n* = 2 the condition is simply *b* > 2*c*, independently of *p*, i.e., the classical Hamilton’s rule is valid. This is so because for *n* = 2 our cost sharing model coincides with the classical model of altruistic actions without cost sharing.

#### When is genotype *G*_3_ evolutionarily stable and when is instable?

By symmetry, conditions for the evolutionary stability of *G*_3_ can be obtained from those of *G*_1_ by interchanging 1 and 3 in the subscripts. Thus the primary condition is *n*_3(33)_ > 2*n*_2(23)_, that is,

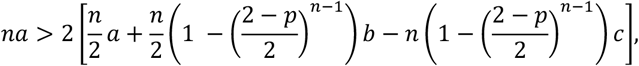

which simply yields 2*c* > *b* whenever *p* > 0. For *p* = 0 we have *n*_3(33)_ = 2*n*_2(23)_, therefore the secondary condition

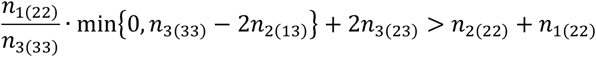

should be applied. After some calculus we get that this is also equivalent to 2*c* > *b*.

Next, we are interested in the *instability* of *G*_3_. Though it is true that reversing the inequality in the primary condition of stability is sufficient for instability, this is not necessarily so for the secondary condition. Thus all we can say is that the classical Hamilton’s rule *b* > 2*c* implies the instability of selfishness provided altruism is not recessive.

### SI-B: When is the net effect of the altruistic interaction remunerative for the altruist gene [a]?

Firstly, let us compute the average benefit of the altruist gene [a] from the interaction. All benefits are given in the following Table S1, for all considered genotype-phenotype mappings and population stages.

Since in genotypes *G*_1_ ≡ ([a],[a]) and *G*_2_ ≡ ([a],[A]) there are 2 and 1 of the altruistic allele [a], resp., we obtain the following expression for the average benefit.

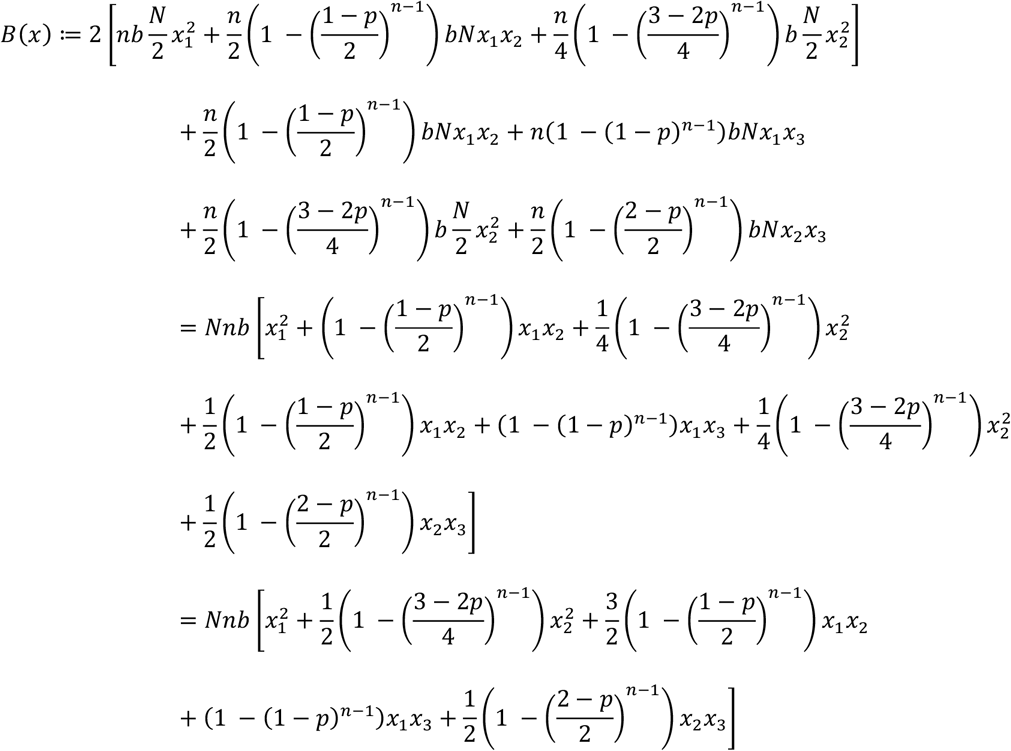

Secondly, let us calculate the average cost of the altruist gene from the interaction. All costs are given in the following Table S2, for all considered genotype-phenotype mappings and population stages.

Similarly, to what we did for the benefit function *B*(*x*), for the average cost we have

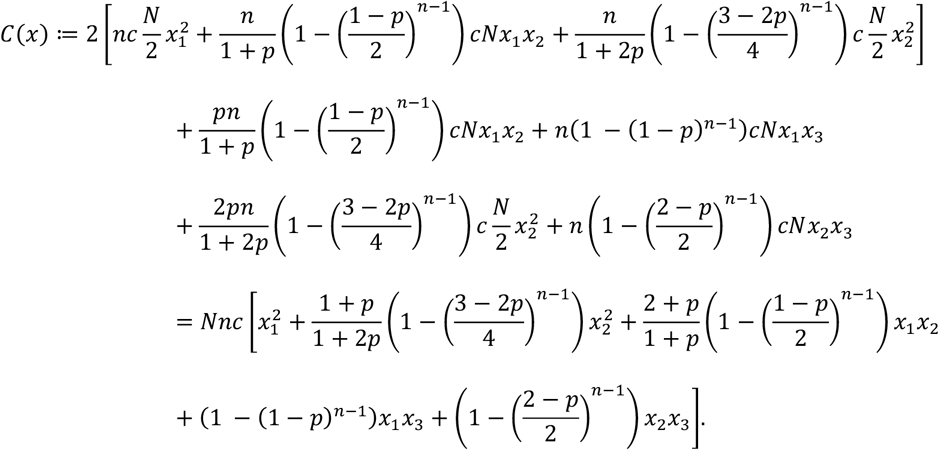

Comparing the benefit and the cost, the function *H*(*x*) = *B*(*x*) − *C*(*x*) shows the total effect of the altruistic interaction for the altruistic gene [a].

For the sake of brevity, we introduce

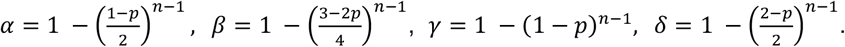

These quantities are positive for *p* > 0. Then

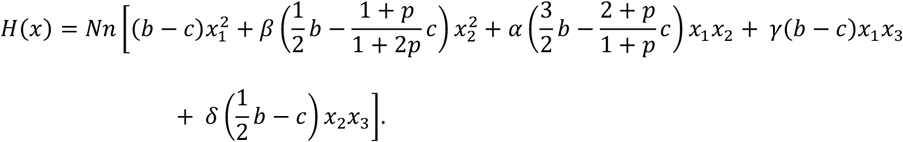

This quadratic form is positive (nonnegative) in the positive octant *x* ≥ 0, if all of its coefficients are nonnegative, i.e., the following conditions are met:

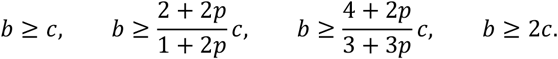

The most stringent bound is *b* ≥ 2*c*; it is sufficient to require this whenever *p* > 0. It is also necessary, because if *x*_1_ = 0 and 0 < *x*_2_ ≪ *x*_3_ (thus *x*_3_ is very close to 1, and *x*_2_ is infinitesimally small), then the dominant term in *H*(*x*) is the one containing *x*_2_*x*_3_. One can easily see that this is also the case for *p* = 0, where γ = δ = 0, and

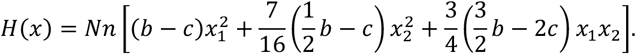

The sufficiency of the condition *b* ≥ 2*c* is obvious, and the necessity also becomes transparent when *x*_1_ = 0 < *x*_2_.

**Table S-1.** The altruistic benefit of altruism in different families.

| Genotypes of parents | Average number of couples | Average number of saved family members, for genotypes $G_1, G_2, G_3$ | | |
| --- | --- | --- | --- | --- |
| | | $G_1 \equiv ([a],[a])$<br>altruistic | $G_2 \equiv ([a],[A])$<br>mixed phenotype $p$ | $G_3 \equiv ([A],[A])$,<br>selfish |
| $G_1 \times G_1$ | $\frac{N}{2} x_1^2$ | $n_{1(11)} = nb$ | $n_{2(11)} = 0$ | |
| $G_1 \times G_2$ | $Nx_1 x_2$ | $\frac{n}{2} \left( 1 - \left( \frac{1-p}{2} \right)^{n-1} \right) b$ | $\frac{n}{2} \left( 1 - \left( \frac{1-p}{2} \right)^{n-1} \right) b$ | |
| $G_1 \times G_3$ | $Nx_1 x_3$ | $n_{1(13)} = 0$ | $n(1 - (1-p)^{n-1})b$ | |
| $G_2 \times G_2$ | $\frac{N}{2} x_2^2$ | $\frac{n}{4} \left( 1 - \left( \frac{3-2p}{4} \right)^{n-1} \right) b$ | $\frac{n}{2} \left( 1 - \left( \frac{3-2p}{4} \right)^{n-1} \right) b$ | |
| $G_2 \times G_3$ | $Nx_2 x_3$ | $n_{1(23)} = 0$ | $\frac{n}{2} \left( 1 - \left( \frac{2-p}{2} \right)^{n-1} \right) b$ | |
| $G_3 \times G_3$ | $\frac{N}{2} x_3^2$ | $n_{1(33)} = 0$ | $n_{2(33)} = 0$ | |

**Table S-2.** The altruistic cost of altruism in different families.

| Genotypes of parents | Average number of couples | Average number of saved family members, for genotypes $G_1$, $G_2$, $G_3$ | | |
| --- | --- | --- | --- | --- |
| | | $G_1 \equiv ([a],[a])$<br>altruistic | $G_2 \equiv ([a],[A])$<br>mixed phenotype $p$ | $G_3 \equiv ([A],[A])$<br>selfish |
| $G_1 \times G_1$ | $\frac{N}{2}x_1^2$ | $n_{1(11)} = -nc$ | $n_{2(11)} = 0$ | |
| $G_1 \times G_2$ | $Nx_1x_2$ | $-\frac{n}{1+p}\left(1 - \left(\frac{1-p}{2}\right)^{n-1}\right)c$ | $-\frac{pn}{1+p}\left(1 - \left(\frac{1-p}{2}\right)^{n-1}\right)c$ | |
| $G_1 \times G_3$ | $Nx_1x_3$ | $n_{1(13)} = 0$ | $-n(1 - (1-p)^{n-1})c$ | |
| $G_2 \times G_2$ | $\frac{N}{2}x_2^2$ | $-\frac{n}{1+2p}\left(1 - \left(\frac{3-2p}{4}\right)^{n-1}\right)c$ | $-\frac{2pn}{1+2p}\left(1 - \left(\frac{3-2p}{4}\right)^{n-1}\right)c$ | |
| $G_2 \times G_3$ | $Nx_2x_3$ | $n_{1(23)} = 0$ | $-n\left(1 - \left(\frac{2-p}{2}\right)^{n-1}\right)c$ | |
| $G_3 \times G_3$ | $\frac{N}{2}x_3^2$ | $n_{1(33)} = 0$ | $n_{2(33)} = 0$ | |

**SI: References**

## Footnotes

1 We note that for haploid sexual populations, where the selection takes place on haploid individuals (gametophytes), the frequency change of haploid genotypes in time, gives the evolutionary change (e.g. [27]), when during the short diploid life period each individual (sporophyte) has the same fecundity and survival rate.

